# Diurnal differences in intracellular replication within splenic macrophages correlates with the outcome of pneumococcal infection

**DOI:** 10.1101/2022.01.30.478422

**Authors:** Ryan G. Hames, Zydrune Jasiunaite, Giuseppe Ercoli, Joseph J. Wanford, David Carreno, Kornelis Straatman, Luisa Martinez-Pomares, Hasan Yesilkaya, Sarah Glenn, E. Richard Moxon, Peter W. Andrew, Charalambos P. Kyriacou, Marco R. Oggioni

## Abstract

Circadian rhythms affect the progression and severity of bacterial infections including those caused by *Streptococcus pneumoniae*, but the mechanisms responsible for this phenomenon remain largely elusive. Following advances in our understanding of the role of replication of *S. pneumoniae* within a specific subset of splenic macrophages, we sought to investigate events within the spleen that correlate with differential outcomes of invasive pneumococcal infection. Utilising murine invasive pneumococcal disease (IPD) models, here we report that infection during the murine active phase (zeitgeber time; 15h after start of light cycle, 3h after start of dark cycle) resulted in significantly faster onset of moderate septicaemia compared to rest phase (zeitgeber time 3; 3h after start of light cycle) infection. These findings correlated with significantly higher pneumococcal burden within the spleen of active phase-infected mice at early time points compared to rest phase-infected mice. Whole-section confocal microscopy analysis of these spleens revealed that the number of pneumococci is significantly higher exclusively within marginal zone metallophilic macrophages (MMMs), known to allow intracellular pneumococcal replication as a prerequisite step to the onset of septicaemia. Pneumococcal clusters within MMMs were more abundant and increased in size in active phase-infected mice compared to those in rest phase-infected mice which decreased in size over time and were present in a lower percentage of MMMs. This phenomenon preceded significantly higher levels of bacteraemia alongside serum IL-6 and TNF-α concentrations in active phase-infected mice following re-seeding of pneumococci into the blood. In summary, these data link the difference in susceptibility to invasive pneumococcal infection to variation in the ability of MMMs to successfully control and digest phagocytosed bacteria.

**Author summary:** Circadian rhythms are present within the majority of multicellular organisms and influence almost all aspects of our physiology. As such, circadian rhythm disorders have been shown to result in an increased susceptibility to certain diseases. The effects of host circadian rhythm have been also mirrored in rodent studies, with the outcome of *Streptococcus pneumoniae* infection being dependent on the time of challenge. Whilst studies into the functional effects of circadian rhythm on the host immune system are present, knowledge of how these contribute to the control of invasive *S. pneumoniae* infection are lacking, especially considering the recent breakthrough in understanding the stages of pneumococcal pathogenesis. We show here that mice infected with *S. pneumoniae* during their active phase developed septicaemia quicker than those infected during their rest phase. We demonstrate that this is likely due to increased replication of pneumococci specifically within a subset of splenic macrophages, which subsequently results in increased numbers of pneumococci in the blood and higher levels of pro-inflammatory cytokines which result in septicaemia. These data provide novel insights into how circadian rhythm influences the immune functionality of the spleen, and how the regulation of function of one macrophage subtype can significantly alter the course of infection.

## Introduction

Circadian clocks have evolved so that organisms can anticipate the relentless changes of light and dark, and hot and cold that are fundamental features of life on a rotating planet. Animals, plants, fungi and certain bacteria predict these environmental changes by regulating their molecular, biochemical, physiological and behavioural cycles [1]. These rhythms include host immune responses to a variety of diseases and bacterial infections including those caused by *Streptococcus pneumoniae* – the leading causative agent of death by communicable disease, despite the availability of effective treatment and vaccines [2-7]. Using highly reliable murine infection models [8], it has been shown that infection with *S. pneumoniae* evidenced a strong periodicity in immune response and outcome of infection, with survival time and time-to-onset of severe bacteraemia being significantly influenced by the time-of-day of infection, and infection during the active phase often resulting in the most favourable outcome [6, 9-11]. However, the immunological mechanisms inducing these diurnal differences, especially with regards to IPD, remain largely undefined.

Recent findings have advanced the understanding of the early stages of IPD at the organ level in murine [12], porcine [13], non-primate and human models [14]. Following intravenous (IV) infection of mice with *S. pneumoniae*, bacteraemia is cleared over a matter of hours primarily via splenic macrophages [12, 13, 15]. However during this time, pneumococci are phagocytosed and replicate to form clusters within two subsets of splenic macrophages: the red pulp macrophages (RPMs) expressing F4/80 and marginal zone metallophilic macrophages (MMMs) expressing Siglec-1 (CD169) [12-14]. While pneumococcal clusters within RPMs are effectively cleared following an influx of neutrophils to the spleen [12, 16], intracellular replication within MMMs precedes the cell lysis which results in the recurrence of bacteraemia and ultimately the onset of fatal septicaemia [12-14]. This stage of pathogenesis is an essential prerequisite for septicaemia as the prevention of pneumococcal uptake by MMMs results in a lack of re-emerging bacteraemia and sepsis, thereby enabling survival of the mice [12, 14].

Given that the spleen and its tissue-resident macrophages play a critical role in the control of pneumococcal infection and the onset of IPD, and that periodicity in response to pneumococcal infection has been recorded previously, we hypothesised that innate immune events involving splenic MMMs would correlate with a diurnal variation in susceptibility to IPD. To this end, we utilised a murine IPD model to show that mice infected during their active phase develop septicaemia faster than those infected during their rest phase. We show here that this ultimately correlates with larger and more numerous clusters of pneumococci within MMMs which precedes increased systemic pro-inflammatory cytokine concentrations of active phase-infected mice.

## Materials and Methods

### Bacterial strains and culture conditions

*S. pneumoniae* serotype 2 strain D39 [17] was plated on brain heart infusion (BHI, Thermo Scientific, MA, USA) agar plates supplemented with 3% v/v defibrinated horse blood (Thermo Scientific) and subsequently grown in BHI broth at 37°C to OD600_nm_=0.3. Cultures were aliquotted and stored at - 80°C in BHI broth supplemented with 10% v/v glycerol (Thermo Scientific) until use. The number of colony forming units (CFU) per mL was calculated from one of the frozen aliquots in the batch before use.

### Balb/cAnN and CD1 mouse strains

Naïve 6-8 week old specific pathogen-free Balb/cAnN or CD1 female mice were sourced from the in-house breeding core facility at the University of Leicester Pre-clinical Research Facility (PRF), or from Charles River (Bristol, UK). Mice were randomised by PRF technicians and housed in groups of 3-5 in individually ventilated cages and given food/water *ad libitum*. Mice were consistently exposed to the PRF’s 12h:12h light:dark lighting schedule (lights on from 07:00-19:00) for at least two weeks prior to any experimental procedures. All animal experiments were performed in accordance with the United Kingdom Home Office project license (P7B01C07A/PP0757060) and were approved by the University of Leicester Ethics Committee.

### Invasive pneumococcal disease mouse model

Mice were infected IV with 100µL of 1×10^6^ CFU of *S. pneumoniae* strain D39 diluted in phosphate-buffered saline (PBS; Thermo Scientific) into the lateral tail vein [15, 18]. Infections were carried out either during the murine rest phase at 10am (zeitgeber time 3; 3 hours after lights on) or during the murine active phase at 10pm (zeitgeber time 15; 15 hours after lights on – 3 hours after lights off). For infections occurring during the dark cycle in the active phase, ambient lighting was kept to a minimum. To determine the time to onset of moderate septicaemia, mice were checked frequently every ∼6h and scored based on the signs of piloerection, hunched posture and lethargic behaviour. Mice were subsequently culled at the humane endpoint, defined as the point of onset of signs of disease consistent with the upper limits of moderate septicaemia (lethargic behaviour combined with significant piloerection and/or very hunched posture). For all experiments measuring bacterial CFU counts in the blood, blood was retrieved via cardiac puncture form mice under terminal anaesthesia and collected in a microcentrifuge tube containing 50 units of heparin (Sigma, MO, US) to prevent coagulation. Spleens and livers were collected and either mechanically homogenised through a 40 µm cell strainer into BHI for determination of CFU load within the organ, or alternatively flash frozen in optimal cutting matrix (OCT, CellPath, Newtown, Wales) using 2-methylbutane (Thermo Scientific) and dry ice for later microscopy analysis. For “dry” organ CFU enumeration, the number of bacteria in the blood within the organ (170µL blood per gram in the spleen and 360µL blood per gram in the liver) at the time point of retrieval was calculated and subtracted from the organ CFU [19].

### Immunohistochemistry, microscopy and image analysis

Organ samples frozen in OCT were cut into 10 µm sections using a CM1850UV cryostat (Leica Biosystems, Wetzlar, Germany) and adhered to polysine microscopy slides (Thermo Scientific). Samples were fixed using 4% v/v Formaldehyde (Sigma) diluted in PEM buffer [20] and stained via immunohistochemistry using various antibodies for visualisation of macrophages and bacteria (S1 Table). Whole spleen sections from 3 mice per group were analysed using an FV1000 confocal laser scanning microscope (Olympus, Tokyo, Japan) equipped with a 60x UPlanSAPO objective (Olympus) for visual determination of foci counts, and 3D reconstructions were created using Imaris software (Oxford Instruments, Abingdon, UK). Entire sections were scanned with 40x magnification using a Vectra Polaris quantitative slide scanner (Akoya Biosciences, MA, US) for subsequent bacterial co-localisation and cell number analysis. Scanned section image analysis was performed using Fiji (v1.53)[21] and inForm image analysis software (v2.5.1; Akoya Biosciences). For determination of the area of pneumococci associated to each macrophage subtype, regions of interest (ROIs) were manually created in Fiji around macrophage marker and bacterial fluorescence signals by appropriate alteration of the fluorescence intensity threshold for each channel. The pixel area of macrophage signal for each subtype, and the area of bacteria within these ROIs, could then be calculated. For determination of the total number of each macrophage subtype within the samples, and the percentage of these associated with bacteria, inForm image analysis software was utilised. Tissue was segmented into macrophage-positive and negative areas for each subtype and cell areas were segmented based on their nuclear staining to give total numbers of each macrophage subtype in the sample. For further analysis, cells were finally phenotyped based on their positivity for pneumococcus using machine learning that was trained by manual selection of at least 20 pneumococci-positive and 20 pneumococci-negative cells.

### Cytokine ELISAs

Serum prepared from whole mouse blood collected at 4-, 12- and 36h post-infection (PI), alongside homogenates of spleens collected at 4h PI, were analysed for the concentration of TNF-α, IL-6 and IL-1β via ELISA kits (Thermo Fisher, MA, US) as per manufacturer’s instructions. Briefly, sera were collected by allowing whole blood to coagulate at room temperature (RT) before centrifuging at 2,000 x g for 20 minutes at RT, and spleens were weighed and mechanically homogenised into BHI + 10% v/v glycerol through a 40 µm cell strainer. Samples were stored at -80°C until use. 96-well plates were coated with antibodies against mouse TNF-α, IL-6 and IL-1β. Wells were washed using PBS + 0.05% v/v Tween-20 (Sigma), non-specific binding was blocked using ELISA diluent (PBS + Bovine serum albumin (BSA)), and a standard was made using seven 1:2 dilutions of recombinant mouse TNF-α, IL-6 or IL-1β in ELISA diluent. Samples (diluted 1:6), standards and blanks were added to duplicate wells and incubated overnight at 4°C. Wells were washed as before and biotin-conjugated antibodies against mouse TNF-α, IL-6 and IL-1β were added and plates incubated for 1 hour at RT. Wells were again washed before the addition of Streptavidin-HRP and a 30-minute incubation at RT. Wells were washed once more, tetramethylbenzidine substrate solution was added and incubated for 15 minutes at RT, before 2M H_2_SO_4_ stop solution was added and absorbance was read at 450 nm and 570 nm. 570 nm absorbance was subtracted from 450 nm absorbance, averages of duplicates were then calculated before blank absorbance values were subtracted from sample absorbance values. Cytokine levels were determined against standard curves generated using CurveExpert (v1.4; Hyams Development). For spleen samples, cytokine concentrations were normalised against their weight to express concentrations as pg per gram of tissue.

## Results

### Time-of-infection imparts a differential susceptibility to invasive pneumococcal infection

To determine whether mice in our IPD model displayed time-of-day dependent differential susceptibility to invasive infection, CD1 mice were IV infected in their rest phase or active phase with *S. pneumoniae* strain D39. CD1 mice were used due to their propensity to develop septicaemia following invasive pneumococcal infection [12, 14, 15]. Mice infected during their active phase developed moderate septicaemia significantly faster time than mice infected during their rest phase, with over 92% of rest phase-infected mice reaching moderate septicaemia by 48h PI, in comparison with only 28% of active phase-infected mice (Fig 1A). This correlated with earlier signs of disease in active phase-infected mice, with the first mouse displaying signs of disease >12h before rest phase-infected mice (Fig 1B).

**Fig 1.**
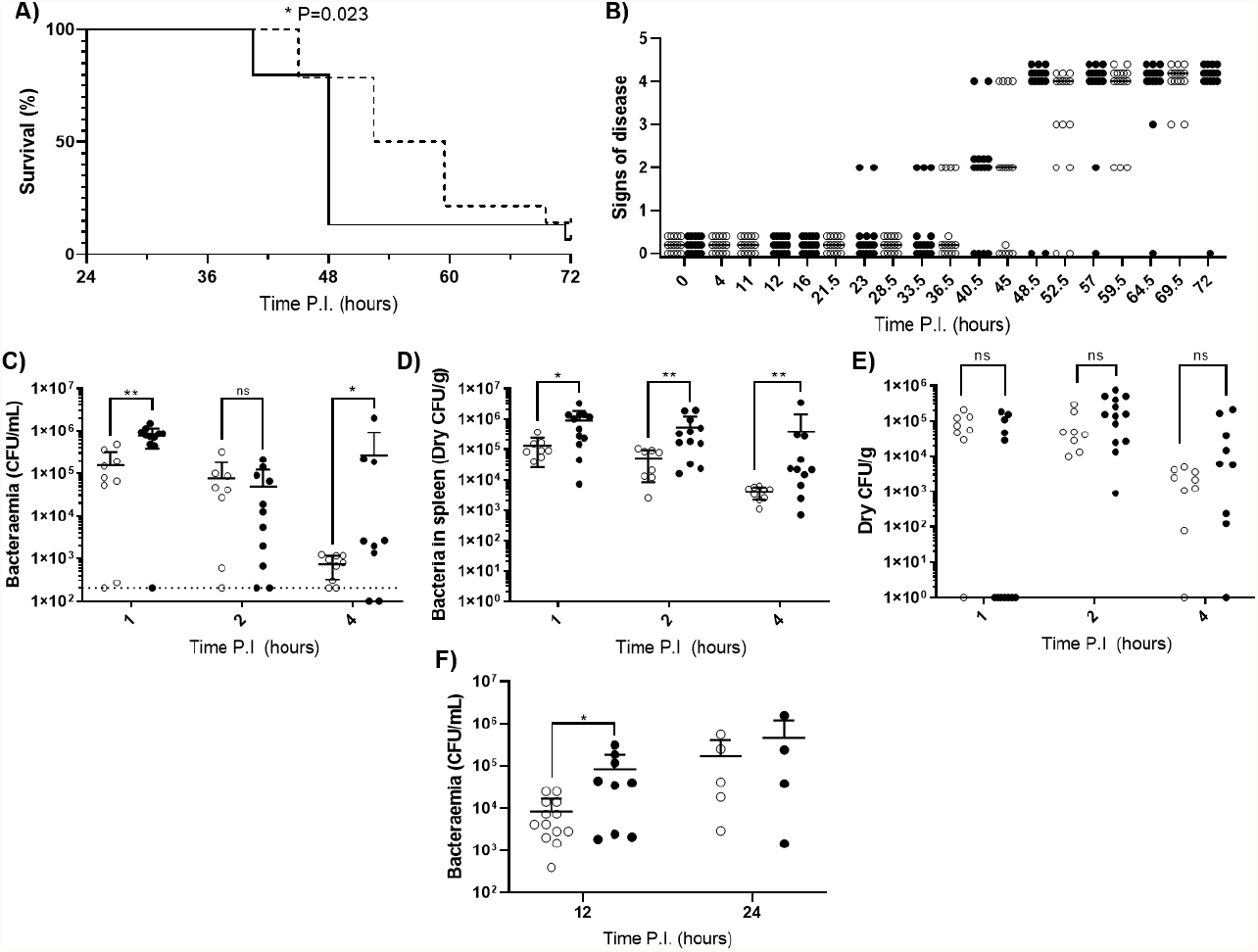
Disease severity and bacterial burdens of blood and organs from mice infected during their rest or active phases. (A) CD1 mice (n=30) were IV infected during their rest (dotted line) or active (solid line) phase and sacrificed once visible clinical signs consistent with moderate septicaemia were reached. Survival curves were compared by Gehan-Breslow-Wilcoxon survival curve test (*; P=0.0232). (B) Mice in (A) were regularly scored for signs of disease equating to 0: normal, 1: hunched, 2: piloerect, 3: hunched + piloerect, 4: lethargic + very hunched or very piloerect. Rest and active phase-infected mice are represented by open and filled points respectively. Blood CFU/mL (C), spleen CFU/g (D), and liver CFU/g (E) bacterial CFU counts were enumerated from mice (n=30) at 1, 2 and 4h PI. Rest and active phase-infected mice are represented by open and filled points respectively. The limit of detection for blood CFU counts is indicated by the horizontal dotted line. Statistical significance was determined at each time point between both groups of mice by Mann-Whitney test (ns; P=0.5438, *; P≤0.05, **; P<0.01). (F) Blood bacteria CFU counts from mice (n=31) were determined at 12 and 24h PI. Rest and active phase-infected mice are represented by open and filled points respectively. Statistical significance was determined at each time point between both groups of mice by Mann-Whitney test (*; P=0.0339).

To identify if the spleen displays diurnal immune responses to pneumococcal infection that could affect septicaemia onset, Balb/cAnN mice were IV infected during their rest phase or active phase and sacrificed at 1, 2 and 4h PI for CFU enumeration of the blood and spleen. The liver was also retrieved due to its role in clearance of other invasive pathogens from the blood [22, 23]. Balb/cAnN mice were utilised here as although they fail to develop septicaemia following invasive infection [15], their inbred phenotype was predicted to reduce variation within groups and emphasise any differences that may occur at early time points. Bacteraemia levels between both groups of mice were different as soon as 1h PI (Fig 1C), indicating potential diurnal differences in the ability of mice to clear bacteria from the blood. Bacteraemia levels were also significantly higher in active phase-infected mice at 4h PI, due to three mice being unable to control the infection. However, dry spleen bacterial burden (excluding blood-borne bacteria within the organ) was significantly greater at all time points in mice infected during the active phase compared to those infected during the rest phase, with splenic bacterial numbers decreasing 30-fold in rest phase-infected mice, compared to only 2-fold in active phase-infected mice (Fig 1D). In comparison, bacterial burden within the liver was comparable between both groups of mice throughout the time course (Fig 1E), highlighting a lack of hepatic circadian-mediated innate immunity against invasive pneumococcal infection within our model.

As the re-seeding of the blood is known to originate from the spleen [12, 14], we also determined the level of bacteraemia immediately following the re-seeding event due to the significantly different numbers of pneumococci recorded within the spleens of both groups of mice. At 12h PI, bacteraemia levels were significantly greater in active phase-infected mice compared to rest phase-infected mice, however no significant difference was recorded at 24h PI (Fig 1F). Taken together, these results confirm that mice display a diurnal susceptibility to IPD that could be correlated to events occurring within the spleen, which in turn influences the magnitude and timing of the re-emergence of pneumococci into the blood.

### Splenic MMMs of active phase-infected mice harbour increased numbers of pneumococci

To determine the reason for the greater bacterial burden in the spleens of mice infecting during their active phase, spleens from three Balb/cAnN mice were taken from each group at 1, 2 and 4h PI and stained, imaged and analysed to quantify bacterial association to the three main subsets of tissue-resident macrophages within the murine spleen: marginal zone metallophilic macrophages (MMMs; CD169+), marginal zone macrophages (MZMs; MARCO+), and red pulp macrophages (RPMs; F4/80+) (Fig S1A-C; S1 Table).

Data show that the area of each splenic macrophage subtype varied within and between samples (S1 Fig), therefore for each sample, bacterial area within each macrophage ROI was expressed as a percentage of the macrophage ROI area. When normalised, RPMs and MZMs yielded the same area of bacteria between rest and active phase-infected mice at all time points, however there was a significantly higher area of pneumococci within MMMs in active phase-infected mice at 4h PI compared to rest phase-infected mice (Fig 2A). To establish whether this increase in pneumococcal association was due to higher numbers of pneumococci per macrophage, or an increased percentage of pneumococci-harbouring MMMs, scans were re-analysed to determine the total number of macrophages belonging to each subtype, and phenotyped based on the presence or absence of pneumococci. These data show that whilst a higher percentage of both MZMs and MMMs were associated with pneumococci at 2h PI in active phase-infected mice compared to rest phase-infected mice, crucially at 4h PI, the same percentage of MMMs were associated with pneumococci between both groups of mice (Fig 2B). In accordance with previous data showing that MMMs and RPMs allow intracellular replication of pneumococci [12, 14], these results demonstrate higher numbers of pneumococci per MMM but not RPM and thus suggest that an increased rate of pneumococcal persistence and replication is occurring exclusively within MMMs in active phase-infected mice.

**Fig 2.**
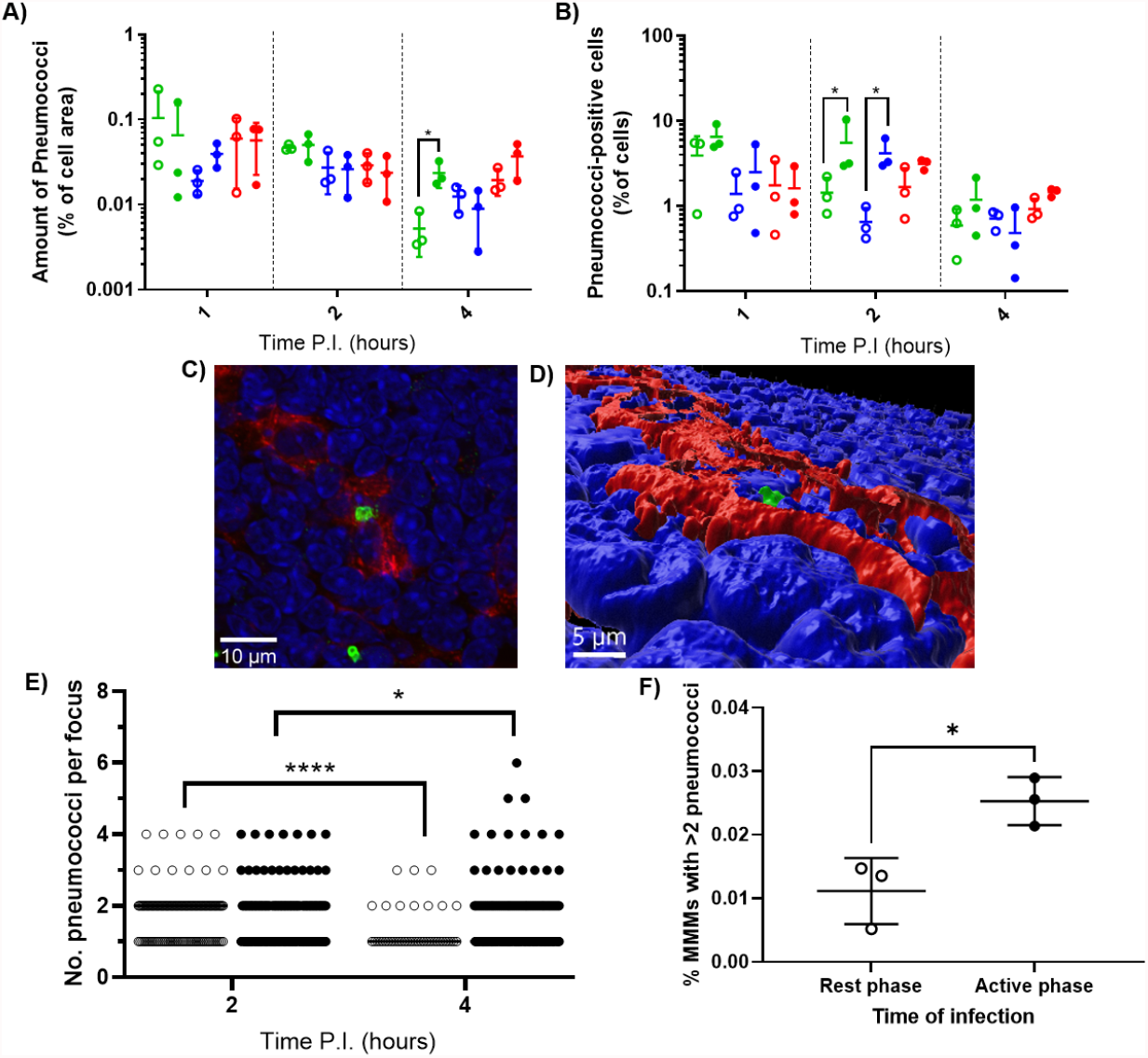
Bacterial localisation and replication within splenic tissue resident macrophages. Spleens from 3 mice per time point per group (n=18) were sectioned and stained for immunohistochemistry to identify MMMs (CD169+), MZMs (MARCO+) and RPMs (F4/80+). (A) Whole sections were scanned and analysed to determine the area of pneumococci associated with each macrophage subtype, which is expressed as a percentage of the macrophage area. MMMs, MZMs and RPMs are represented by green, blue and red points respectively. Rest and active phase-infected mice are represented by open and filled points respectively. Statistical significance was determined at each time point by Mann-Whitney test between both groups of mice for each macrophage subtype (*; P≤0.05). (B) Spleen sections were scanned using a quantitative slide scanner and analysed to determine the percentage of each macrophage subtype that are associated with pneumococci. MMMs, MZMs and RPMs are represented by green, blue and red points respectively. Rest and active phase-infected mice are represented by open and filled points respectively. Statistical significance was determined at each time point by Mann-Whitney test between both groups of mice for each macrophage subtype (*; P≤0.05). (C) Confocal single optical section image of a 6-pneumococci cluster associated with a MMM at 4 hours PI showing cell nuclei (blue), CD169 (red) and pneumococci (green). The scale bar represents 10µm. (D) 3D-reconstruction of (A) showing the pneumococcal cluster (green) within the confines of CD169+ signal (red) verifying intracellularity. Nuclei are shown in blue. The scale bar represents 5µm. (E) 2 and 4h PI spleen samples stained to visualise MMMs were manually analysed using confocal microscopy. The number of intracellular pneumococci within MMMs were recorded. Rest and active phase-infected mice are represented by open and filled points respectively. Statistical significance was determined between time points for each group by Mann-Whitney test (*; P=0.0327, ****; P<0.0001). (F) Whole sections analysed in (E) were re-analysed to determine the number of MMMs within each section. The total number of pneumococcal clusters consisting of 3 or more bacteria was normalised against the total number of MMMs to give the percentage of MMMs that contain replicative foci at 4h PI. Rest and active phase-infected mice are represented by open and filled points respectively. Statistical significance was determined Mann-Whitney test (*; P=0.0189).

To confirm this, three spleen samples per group at 2h and 4h PI were analysed to determine the number of pneumococci per macrophage specifically within infected MMMs. All visible bacteria in each field of view were assessed, and those deemed to be intracellular due to complete localisation within the confines of the cell membrane CD169+ stain were recorded (Fig 2CD). These data reveal the average number of intracellular pneumococci per MMM significantly decreased from 2-4h PI in rest phase-infected mice, whilst increasing in active phase-infected mice (Fig 2E). These results suggest a higher tolerance for pneumococcal persistence and replication within MMMs of active phase-infected mice. To strengthen this hypothesis, when considering the total number of MMMs within these samples, a higher percentage of MMMs were found to contain replicative foci (>2 pneumococci) in active phase-infected mice (Fig 2F), indicating a higher propensity of these macrophages to permit intracellular persistence and replication in active phase-infected mice.

### Active phase-infected mice display increased TNF-α and IL-6 levels only after blood re-seeding

Septicaemia is inherently moderated by cytokines [24, 25]. Therefore, to explore the effect of differential pneumococcal replication within the spleen on serum cytokine levels, and to determine whether the differential phenotype displayed by MMMs is mediated by local cytokine levels, cytokine concentrations in serum and spleen homogenates were analysed for the quantification of cytokine concentrations post-pneumococcal infection. Sera were retrieved from whole blood samples collected at 4, 12 and 36h PI from CD1 mice, and spleens were collected at 4h PI. Samples were analysed by quantitative ELISA to determine the concentration of pro-inflammatory cytokines IL-1β, TNF-α and IL-6 at each time point. In both groups of mice, and for all cytokines tested, the concentration of cytokines in the sera of both groups of mice at 4 and 12h PI were not significantly different, implying that differential levels of cytokines are not produced in response to the same levels of bacteraemia (Fig 3ACE). However, whilst increasing in both groups of mice, TNF-α and IL-6 concentrations in the sera were higher at 36h PI in active phase-infected mice, at around 24h after the onset of bacteraemia resulting from splenic re-seeding (Fig 3AC). IL-1β remained low throughout the time course for both groups of mice (Figure 3E), whilst concentrations of cytokines from uninfected samples were below the limit of detection. These results suggest that the release of bacteria into the blood following differential pneumococcal replication events within the spleen is responsible for differential cytokine production, which in turn reflects the differential time-to-onset of moderate septicaemia. Further, the similar concentrations of cytokines in the spleen at 4h PI indicate that other non-cytokine mediated circadian factors are likely driving the differential phenotypes of MMMs (Fig 3BDF).

**Fig 3.**
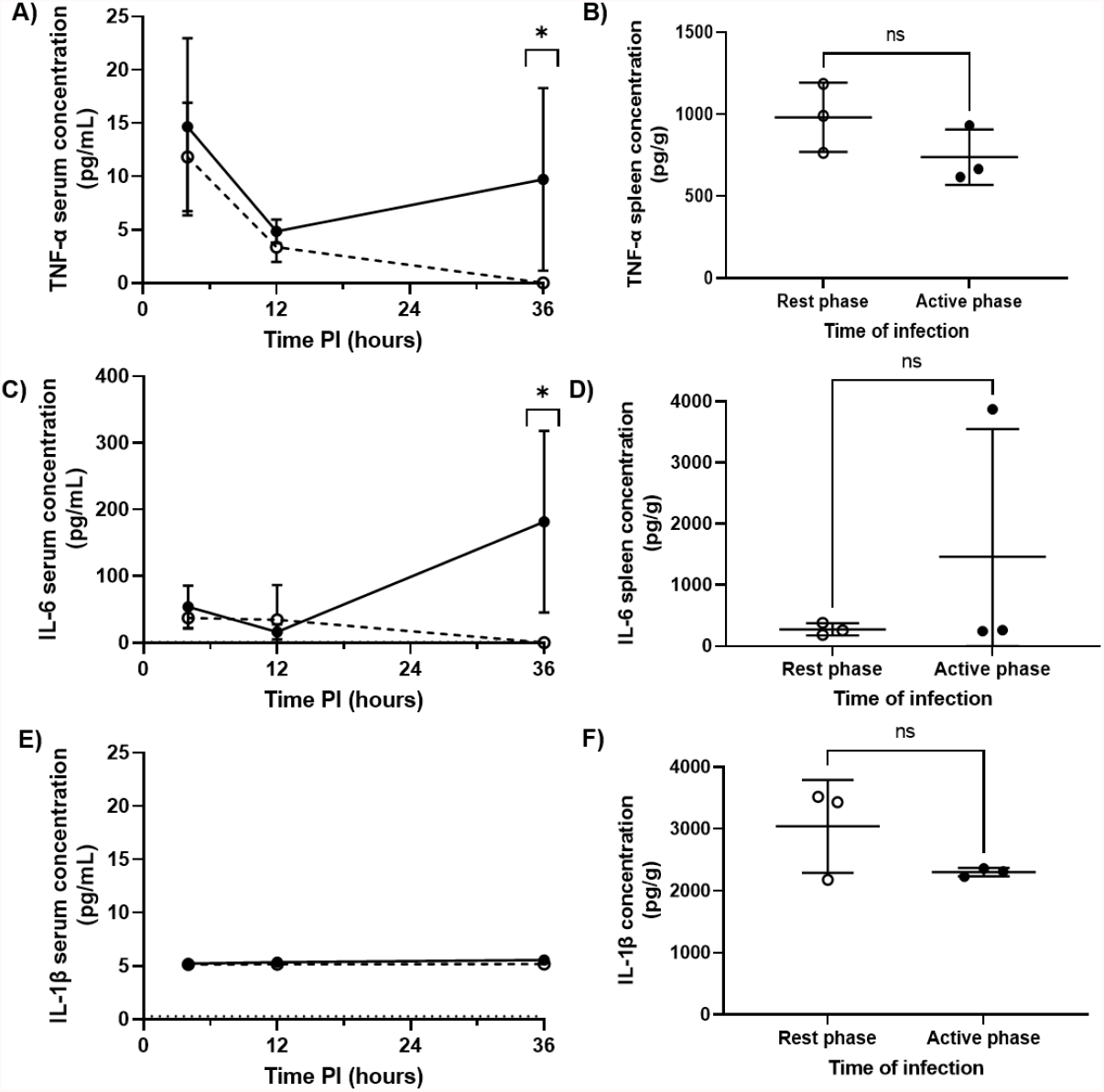
Serum and splenic cytokine concentrations of TNF-α, IL-6 and IL-1β in rest and active phase-infected mice. Serum and spleen homogenates were retrieved from CD1 mice (n=18) at 4h (spleens+serum), 12h (serum) and 36h (serum) PI. Samples were analysed for TNF-α (AB), IL-6 (CD) and IL-1β (EF) concentrations. Rest and active phase-infected mice are represented by open and filled circles respectively. Statistical difference was determined using Mann-Whitney test between both groups of mice at each time point (ns; P>0.05, *; P≤0.05).

## Discussion

Diurnal periodicity in the fate of mice infected with *S. pneumoniae* has been documented [6, 9-11], however the immune mechanisms behind these differences, especially with regards to IPD, remain elusive. Evidence suggests that the spleen plays a pivotal role in IPD onset [12, 26] including when induced by high severity pneumonia [14]. Thus, we explored whether host-pathogen interactions within the spleen display diurnal variation following pneumococcal infection, and if so, whether the differences correlate with the subsequent outcome of IPD. Utilising *in vivo* murine IPD models in which infections began during the rest phase or active phase and immunohistochemical analysis of spleens, we found that increased splenic bacterial load and a higher incidence and size of bacterial foci within MMMs in mice infected during their active phase correlates with a more pronounced re-emergence of bacteria into the blood, and an increase in systemic pro-inflammatory cytokine levels resulting in a shorter time-to-onset of moderate septicaemia.

Mammalian circadian rhythms modulate various functions of the innate immune system in order to coincide with the likelihood of host exposure to pathogens, often during the organism’s active phase [27, 28]. Our demonstration of a significant difference in the time-to-onset of septicaemia in mice infected in opposite phases of the circadian cycle reflects this notion. Interestingly however, we show that utilising an IPD murine model, mice infected during their active phase reach moderate septicaemia significantly faster than mice infected during their rest phase. Whilst other studies reveal a more favourable outcome when pneumococcal infection is initiated during the active phase, these models employed subcutaneous, intraperitoneal and intranasal routes of infection, all of which involve host immune interactions prior to the seeding of the blood and onset of IPD [6, 9, 11]. Indeed, circadian-driven fluctuations in CXCL5 expression by pulmonary epithelial club cells results in increased pulmonary inflammation and neutrophil infiltration contributing to significantly lower pneumococcal bacteraemia in mice infected during their active phase [6]. In the context of intraperitoneal infection, control of the bacterial challenge prior to the onset of bacteraemia is first initiated by peritoneal macrophages, which themselves display significantly increased phagocytic index during the late-active phase compared to other points of the circadian cycle [29]. Therefore, the overall outcome of infection is likely to depend on the model and route of infection; challenge in a natural host and with a natural route of infection will likely involve multiple factors that display diurnal oscillation, with some carrying more weight than others with regards to the progression towards disease or remission. Nonetheless, it is enticing that our IPD model, which avoids confounding immune interactions prior to the onset of IPD and emphasises tissue macrophage activity in the spleen, highlights potential influences of circadian rhythm on the host-pathogen interactions between bacteria and splenic tissue-resident macrophages.

We next determined that mice infected during their active phase presented significantly higher splenic bacterial burdens at all time points, with the largest difference observed at 4h PI. Immunohistochemical analysis of spleen sections taken at this time point from both groups of mice revealed pneumococcal clusters within MMMs and RPMs due to intracellular replication, as previously reported [12-14]. Whilst RPMs possess the ability to allow pneumococcal replication [12], pneumococci within this subset of macrophages appears to be effectively controlled by neutrophil influx into the splenic red pulp [12, 16], and a difference in bacterial area within these cells was not recorded between both groups of mice in our study. In our samples, pneumococcal clusters appearing within the MMMs of active phase-infected mice increased significantly in size over time and were present in a higher percentage of MMMs compared to rest phase-infected mice in which cluster size also decreased over time. When taken alongside similar bacterial area within RPMs and MZMs in both groups of mice, and a lack of any known circadian rhythms in *S. pneumoniae*, this finding accounts for the increased splenic bacterial burden we recorded in active phase-infected mice. MMMs, due to their antigen presenting properties and proximity T- and B-cells in murine white pulp follicles, are known to allow the replication of viral particles to increase the quantity of antigen for presentation, and thus promote adaptive immune responses [30]. It is therefore tempting to suggest that the circadian-dependent regulation of the host immune response results in an increased ability of MMMs to perform this function, therefore permitting the formation of larger and more numerous pneumococcal foci. If true, the diurnal modulation of MMM function described herein could hold implications for further understanding the differential efficacy recorded when some vaccines are administered at alternate points of the circadian cycle [31, 32], in-keeping with current knowledge of circadian fluctuations in T-cell function also impacting vaccine efficacy [33]. It is still unclear, however, whether the intracellular replication of pneumococci within MMMs arises due to a hijacking of this cellular response or is merely an exploitation of MMMs decreased role in pneumococcal clearance compared to neighbouring MZMs or RPMs [16, 34, 35]. Although circadian effects on macrophage phagocytosis have been readily studied, the effects on intracellular killing and digestion mechanisms have not been extensively elucidated and thus warrant further study [36].

Finally, as circulating levels of pro-inflammatory cytokines and their expression by macrophages in response to infection are known to display diurnal fluctuation [37, 38], we explored whether the shorter time to onset of septicaemia in active phase-infected mice could also be mediated by diurnal cytokine expression acting independently of the differential replication phenotype that we report within the spleen. We found that the serum concentration of IL-6 and TNF-α, two pro-inflammatory cytokines regularly associated with sepsis [39-41], differed between both groups of mice only after the re-emergence of bacteria from the spleen. Therefore, it can be reasonably proposed that replication within, and/or release of pneumococci from, MMMs is an essential pre-requisite for the generation of differential cytokine levels. The increase in bacteraemia observed at 12h PI in active phase-infected mice could itself result in increased cytokine production, however pneumococci are also known to induce pyroptotic and necroptotic cell death in various tissue resident macrophages [42, 43]. The necroptosis and pyroptosis pathways are known to result in the production of pro-inflammatory cytokines including TNF-α and IL-6 either through direct signalling or indirectly through the release of intracellular damage-associated molecular patters (DAMPs) [44-46]. It is therefore plausible that an increase in the number of persistent pneumococcal clusters within MMMs of active phase-infected mice results in increased numbers of these cells undergoing pyroptotic or necroptotic cell death, and a resultant increase in pro-inflammatory cytokine production, although further study is required to confirm this hypothesis. Further, local levels of these cytokines within the spleen were comparable between both sets of mice at 4h PI suggesting that the diurnal phenotype of MMMs is likely due to cell-autonomous circadian pathways, and not mediated by localised or systemic differences in cytokine levels [37, 47].

The results herein indicate a correlation between the propensity of splenic MMMs to allow intracellular persistence and replication of pneumococci, and the overall outcome of IPD. Further, we build on previous findings demonstrating that pneumococcal-MMM interaction is the stage of pathogenesis that determines whether invasive pneumococcal infection is controlled, by presenting evidence to show that this step is also the rate-limiting stage of pneumococcal pathogenesis therefore mediating both disease progression rate and severity.

## Acknowledgements

Authors thank the staff of the Leicester Preclinical Research Facility for support with the mouse experiments and the University of Leicester Advanced Imaging Facility (RRID:SCR_020967) for use of microscope equipment. The work was in part supported by a collaboration agreement with the University of Oxford and grants from the MRC MR/ M003078/1 and BBSRC BB/S507052/1 to MRO. ZJ is funded by BBSRC BB/S507052/1.

## Contributors

All authors read and approved the final version of the manuscript. RGH, DC, JJW, GE and ZJ performed experiments and wrote the manuscript, KS supervised microscopy, LMP provided guidance on immunology, SG, HY and PWA, supervised the mouse experiments, and ERM, CPK and MRO did the project planning, contributed funding and verified the underlying data.

## Supporting information

**S1 Fig.**
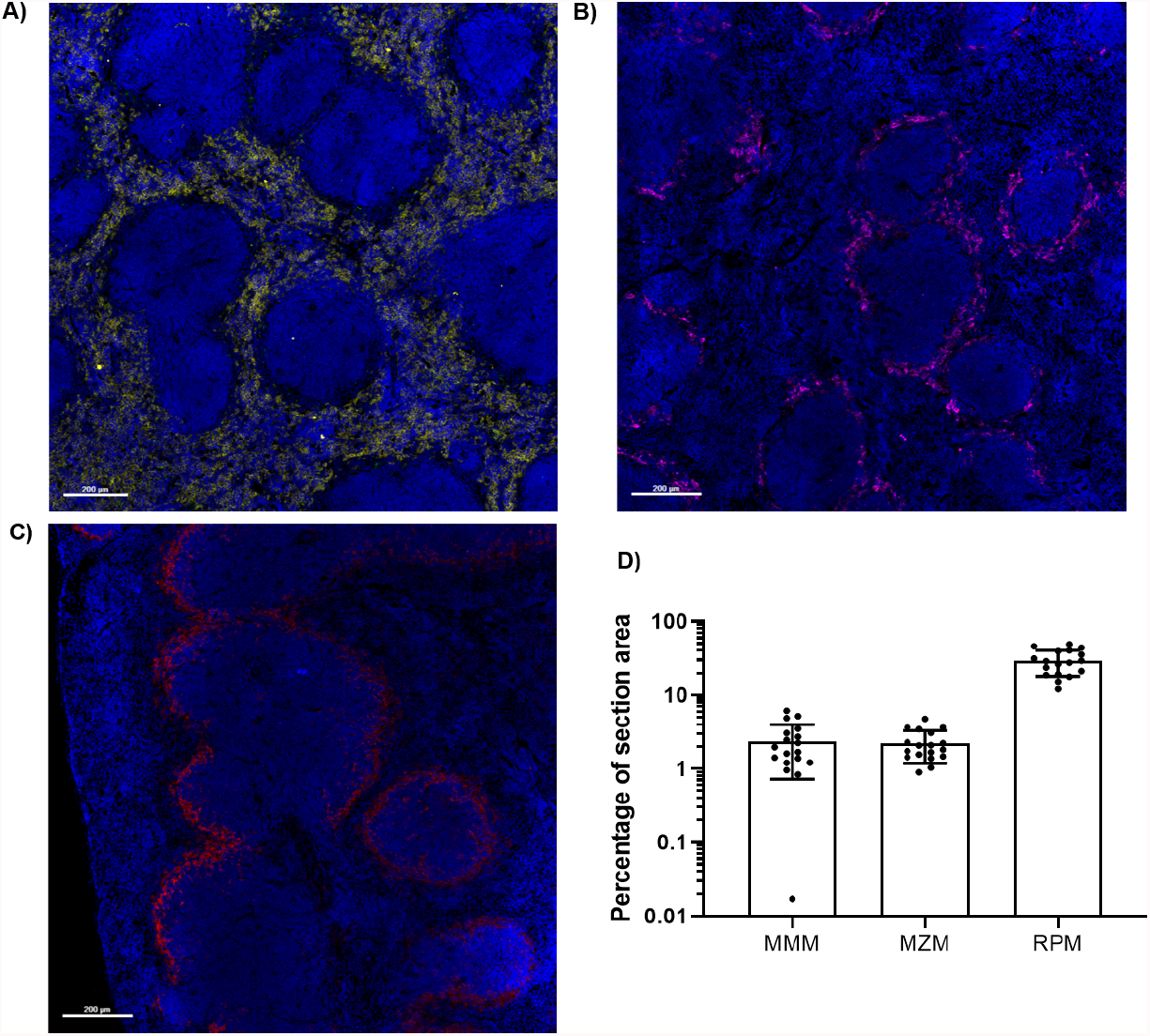
Distribution of macrophage subsets in the murine spleen. Murine spleens were stained for immunohistochemistry and visualised via scanning confocal microscopy to show **(A)** RPMs (F4/80+; yellow), **(B)** MZMs (MARCO+; magenta) and **(C)** MMMs (CD169+; red). Nuclei are stained using DAPI and appear blue. **D)** Images of spleens analysed in Figure 4 (n=54), were analysed to determine the pixel area of MMMs (CD169+), MZMs (MARCO+) and RPMs (F4/80+) as a percentage of the entire section area. Dots represent each analysed spleen. Statistical significance was determined using Dunn’s multiple comparison test (ns; P>0.9999, ****; P<0.0001).

**S1 Table.** Antibodies used for immunohistochemistry.

| Antibody | Specificity | Clone | Conjugated | Catalogue | Supplier |
| --- | --- | --- | --- | --- | --- |
| Purified anti-mouse CD169 (Siglec-1) | MMMs | 3D6.112 | No | 142402 | BioLegend |
| Anti-mouse/human/rat F/80 | RPMs | BM8 | No | 14-4801-82 | Thermo Fisher Scientific |
| Anti-mouse MARCO | MZMs | ED31 | No | MCA1849 | Bio-Rad |
| Anti-type 2 pneumococcal capsule | Type 2 pneumococci | - | No | 16745 | Statens Serum Institut |
| Donkey anti-Rabbit IgG (H+L) | Rabbit IgG (secondary Ab) |  | Alexa Fluor 488 | A-21206 | Thermo Fisher Scientific |
| Goat anti-Rat IgG (H+L) | Rat IgG (secondary Ab) | - | Alexa Fluor 568 | A-11077 | Thermo Fisher Scientific |

